# Deep embeddings to comprehend and visualize microbiome protein space

**DOI:** 10.1101/2021.07.21.452490

**Authors:** Krzysztof Odrzywolek, Zuzanna Karwowska, Jan Majta, Aleksander Byrski, Kaja Milanowska-Zabel, Tomasz Kosciolek

## Abstract

Understanding the function of microbial proteins is essential to reveal the clinical potential of the microbiome. The application of high-throughput sequencing technologies allows for fast and increasingly cheaper acquisition of data from microbial communities. However, many of the inferred protein sequences are novel and not catalogued, hence the possibility of predicting their function through conventional homology-based approaches is limited. Here, we leverage a deep-learning-based representation of proteins to assess its utility in alignment-free analysis of microbial proteins. We trained a language model on the Unified Human Gastrointestinal Protein catalogue and validated the resulting protein representation on the bacterial part of the SwissProt database. Finally, we present a use case on proteins involved in SCFA metabolism. Results indicate that the deep learning model manages to accurately represent features related to protein structure and function, allowing for alignment-free protein analyses. Technologies that contextualize metagenomic data are a promising direction to deeply understand the microbiome.

## Introduction

In just over a decade, a substantial body of evidence linked gut microbiome dysbiosis with diseases ranging from obesity^1^, inflammatory bowel disease^2–4^, diabetes^5,6^, cancer^7,8^, depression^9^ and other psychiatric disorders^10,11^. It shows the profound impact of the microbiome on human health and is a testament to rapid technological progress in sequencing technologies. Since the mid-2000s, the bulk of our insight into the role of the microbiome came from high-throughput and cost-effective 16S rRNA marker gene sequencing experiments that allow for taxonomic discrimination between microorganisms. Though informative, microbiome analysis based solely on taxonomy is prone to bias, due to incomplete reference databases and does not provide detailed information about microbiome function^12^. One of the areas of high interest and relevance is our ability to deduce the gene function from sequence, as it provides more insight into the microbiome’s role in human health. Functional analysis of microbiome data can be performed based on high-throughput, large-scale shotgun metagenomics and other multi-omics experiments that are now becoming accessible for large-scale studies. Gene sequence fragments generated during a shotgun sequencing experiment can be functionally annotated, using homology-based tools such as BLAST or HMMER that search fragments of sequences against reference databases such as Pfam or Gene Ontology (GO)^13^. Similarly to 16S sequencing, functional assignment can be biased, due to incomplete reference databases; so far, only up to 50% of all microbial protein sequences may be annotated^14^. Despite remarkable progress in the last decades, developing precise methods for function prediction is still a major challenge in bioinformatics (see CAFA^15^ initiative). The volume of metagenomic data is making the problem even more difficult to deal with. Thus, introducing an *in silico* method to help assign protein functions could prove highly beneficial for realizing the full potential behind metagenomics and multi-omics.

Deep learning is a proven technique for dealing with intricate problems and has been shown to work well for tasks like speech recognition, natural language processing, or image classification^16^. Recently, it has been successfully employed for analyzing biological sequences, like genomes, proteomes^17^ or metagenomes^18^. Perhaps the best-known example of the use of deep learning in biology was the protein structure prediction problem. DeepMind’s AlphaFold models^19–21^ won the last two Critical Assessment of protein Structure Prediction (CASP) challenges - CASP13^22^ and CASP14^23^, bringing a seismic shift to this decades-old field. The main reason for the notable success of Deep Neural Networks in these areas of biology is their ability to process massive amounts of data, even unlabeled, and extract meaningful patterns from them. Deep learning can leverage the exponential growth of data available in biological databases, which may be limiting for traditional methods. The capability to learn from unlabeled data is particularly valuable due to the constantly increasing gap between the number of unlabeled and labeled protein sequences (https://www.uniprot.org/statistics/TrEMBL).

So far, deep learning methods in protein bioinformatics were employed in two ways: to directly annotate the sequence (supervised learning) or to create a representation of a protein (for example, a sequence embedding using self-supervised learning). Annotation using deep learning is a natural extension of traditional methods, which aim to assign a label to a newly sequenced protein. The label is usually connected to an entry from a database of choice and may belong to curated ontologies (e.g., GO terms^24^) or classification schemes (e.g., EC numbers^25^). Accordingly, studies in the last decade show that deep learning can successfully predict EC numbers^26,27^, GO terms^28–33^, Pfam families^34,35^, or multiple labels at once^36^. However, the labeled proteins are not only in shortage, limiting the potential of deep learning, but also skewed towards model organisms, which may result in biased models.

To overcome these obstacles, more recent approaches use massive unlabeled datasets (UniParc, BFD, Pfam) to train self-supervised models. These models analyze raw amino acid sequences in an alignment-free fashion to learn statistical representations of a protein. The representation can then be effectively used for downstream analyses and predictions of, e.g. secondary or tertiary structure, protein stability, contact map^37,38^, protein function^39,40^, localization^41,42^, variant effect^43^, protein engineering^43,44^, remote homology detection^37^ and more. Moreover, deep-learning-based methods can be used to analyze proteins that do not resemble any catalogued proteins, which is particularly useful in the case of the under-annotated microbiome protein space. Deep-learning-based representations are computationally efficient and accurate, hence they seem appropriate to leverage large amounts of data in high-volume metagenomic studies. Compared to standard bioinformatic tools used to functionally annotate sequences, such as BLAST and HMMER, deep-learning methods require a larger amount of resources, but only during the training phase.

Here, we describe a deep learning approach, based on BiLSTM (Bidirectional Long Short-Term Memory) model, which leverages deep sequence embeddings to understand their potential for solving metagenomic challenges. We trained the model on 20 million microbial proteins from the Unified Human Gastrointestinal Protein (UHGP) catalogue, and then demonstrated the utility of the proposed representations on the Bacterial SwissProt database.

In the first part of this paper, we assessed the type of information encoded in the embedding space. In the second part, we visualized and interpreted the space using Uniform Manifold Approximation and Projection (UMAP)^45^, which allowed for a better interpretation of the evaluation results. As an extension, we built an interactive visualization of the space, which is available at https://protein-explorer.ardigen.com. Finally, we showed the advantages of the embeddings on an example of short-chain fatty acid kinases.

The use of deep protein representations can be beneficial in metagenomic studies, providing advantages over sequence homology-based approaches in terms of computation time and annotation coverage. A deep model can create a global protein space, strongly related to protein function, by making use of unannotated protein sequences in an unsupervised manner. Representing proteins in this space enables their rapid analysis, using a wide range of traditional methods operating on vector spaces and facilitates tasks, such as classification, clustering or semantic search. The model learns abstract patterns that combine, but also go beyond protein sequence and domain architecture. The use of representation space enables to group even sequentially distant proteins into clusters of proteins sharing similar functions. The speed and accuracy of the model is promising in applied settings, such as predicting the mode of action of bacteria and their pathways, new therapeutic design, etc., where the time and ease of use of computational tools may provide accurate interpretations of generated data in a matter of minutes.

## Results

### Alignment-free deep protein embeddings represent structure- and function-related ontologies

Metagenomic data may generate an amount of information on the order of tens of millions of reads, which may be assembled into millions of protein sequences. For traditional sequence homology or profile-based approaches, this amount of data is manageable, but requires significant computing power. For deep learning, on the other hand, such a large amount of data provides an opportunity to be exploited for training and assures a robust representation of analysed sequences.

To build the deep representation, we trained the BiLSTM model on the Unified Human Gastrointestinal Protein catalog (UHGP), which contains 625 million microbial protein sequences clustered with MMseqs2 linclust into 20,239,340 representative sequences at 95% amino acid sequence identity^14,46^. From the trained model, we take a hidden-state vector that acts as a protein representation (see Methods and Fig. 1A).

**Figure 1.**
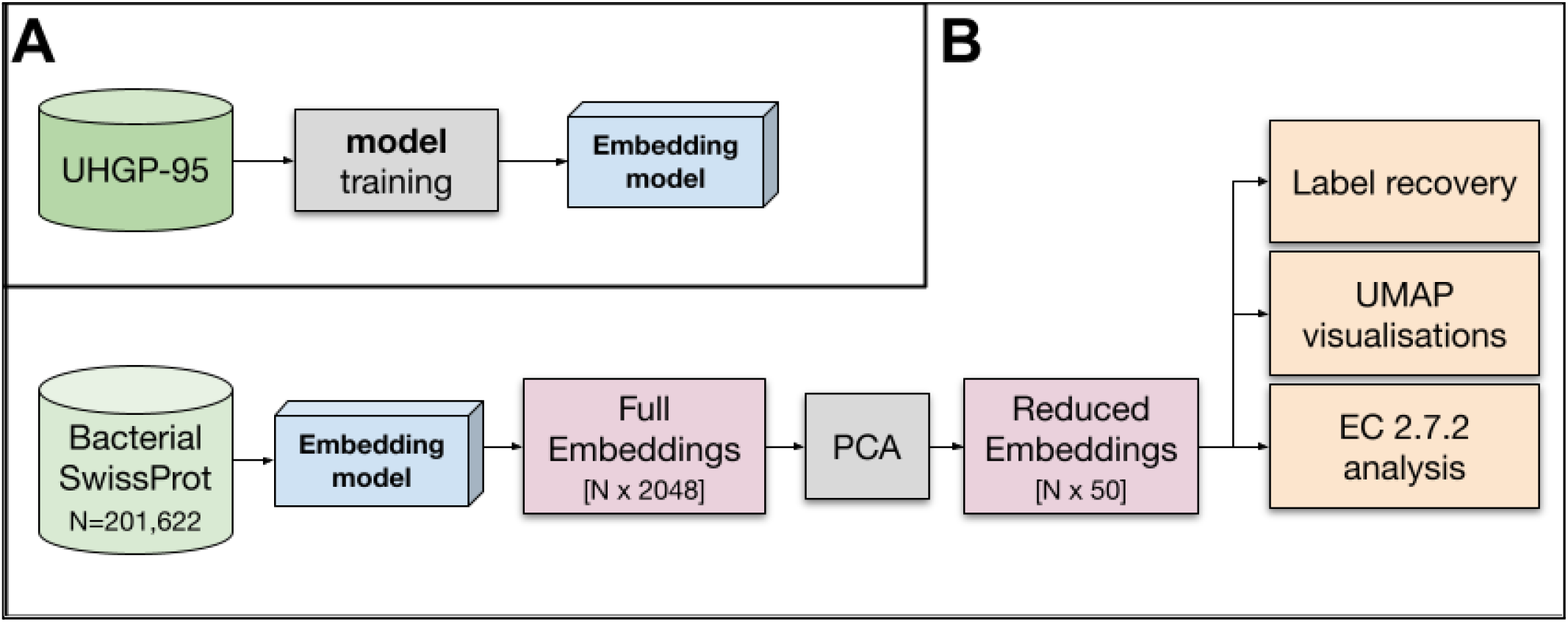
Workflow showing the training of the model and its subsequent use in analyzes.

Our ultimate goal is to produce reliable embeddings for metagenomic data, hence we first validated the model on proteins derived from ten metagenomic samples that were not included in the UHGP catalog (see Methods for details). The model yielded substantially lower Exponential Cross-Entropy (ECE) loss (10.9 ± 0.4) than the analogous model trained on the Pfam database (15.3 ± 0.6). Although ECE loss does not directly measure the quality of obtained embeddings, it was proven that the lower ECE the better the embeddings are in secondary structure and contact predictions^38^. Pfam is a cross-sectional curated dataset built on top of UniProtKB and is limited to identified protein families (~77% of UniProtKB sequences). The UHGP, on the other hand, is a more comprehensive database for gut metagenomic samples which in many cases (up to ~40%) are not represented in protein classification databases (eg. InterPro), and consequently in Pfam^14^. This emphasizes the importance of training an embedding model on a set of proteins consistent with the investigated dataset, i.e. human gut metagenomic proteins. Taken together, this leads to improved model performance.

Although the representation is aimed for metagenomic data, we need proteins with a specified function and origin to validate it. Therefore, for our analysis, we used bacterial proteins from the SwissProt database clustered into 201,622 representatives at 97% sequence identity. SwissProt is a reliable source, linking proteins to many ontologies that enable a multilevel description of sequences (e.g. Table 1). For simplicity, we call this collection of proteins Bacterial SwissProt (see Methods). We generated embeddings for all Bacterial SwissProt sequences using the embedding model and then reduced the 2,048 dimensional vectors with Principal Component Analysis (PCA) to 50 dimensions. Such a representation is used in all our analyzes (Reduced Embeddings in Fig. 1B). Rationale for selected parameters can be found in the Methods section.

**Table 1.** Description of Bacterial SwissProt ontology databases. For the label recovery task, we used a number of ontologies that can be assigned to a protein. These ontologies are based on 3D protein structure (SUPFAM, Gene 3D), domains (Pfam, InterPro), function (GO, KO, EC numbers) or provide information about organism of origin (taxonomy)

| Database | Category | Description | Bacterial SwissProt |  |
| --- | --- | --- | --- | --- |
|  |  |  | #proteins | #classes |
| **SUPFAM** | Structure | SUPFAM associates sequence families from Pfam with SCOP structural families using profile matching to produce sequence superfamilies of known structure. | 147,137 | 989 |
| **GENE 3D** | Structure | GENE 3D contains protein domain assignments for sequences from all of the major sequence databases. Domains are predicted using a library of representative profile HMMs, derived from CATH superfamilies or directly mapped from structures in the CATH database. | 116,919 | 1,173 |
| **InterPro** | Sequence and domain | InterPro brings together 11 protein family databases (CATH-Gene3D, HAMAP, PANTHER, Pfam, PRINTS, ProDom, PROSITE Patterns, PROSITE Profiles, SMART, SUPERFAMILY, and TIGRFAMs). Each database provides a specific signature i.e. position-specific score matrices, hidden Markov models and profiles etc. to increase the sensitivity of protein classification. | 198,677 | 12,244 |
| **KO (KEGG Orthology)** | Function | KO is a database of molecular functions. Each molecular function is represented in terms of a manually defined functional ortholog that together create molecular networks (pathways). Each functional ortholog is defined from experimentally characterized genes and proteins in specific organisms, which are then used to assign orthologous genes in other organisms, based on sequence similarity. | 177,018 | 6,614 |
| **GO (Gene Ontology)** | Function | GO is a controlled terminology that can be used to consistently and structurally identify genes and gene products. The GO terms are organized within a directed acyclic graph (DAG), and each GO term has a described relationship to one or more other terms in the same domain (i.e. biological process, molecular function, or cellular location). | 192,990 | 5,799 |
| **eggNOG** | Function and taxonomy | eggNOG is a database of orthology relationships, gene evolutionary histories and functional annotations. It is built on the concept of OGs (orthologous groups) that are the result of a non-supervised analysis of thousands of genomes and relationships between all their genes. | 162,261 | 15,932 |
| **EC number** | Function | EC numbers are a manually assigned nomenclature that describes enzymes, based on the chemical reactions they catalyse. | 193,198 | 3,005 |
| **Pfam** | Sequence | Pfam is a database of protein families and domains. Each | 120,184 | 5,551 |
|  | and domain | Pfam family has a seed alignment that contains a representative set of sequences for the entry. This alignment is used to build a hidden Markov model profile and the profile is being searched in the sequence database called pfamseq using the HMMER software. |  |  |
| **Taxonomy:**<br>**Order** | Taxonomy | Uniprot uses the NCBI taxonomic database to assign taxonomic identifiers to nucleotide sequences. | 200,536 | 132 |
| **Taxonomy:**<br>**Family** |  |  | 198,996 | 274 |
| **Taxonomy:**<br>**Genus** |  |  | 200,615 | 660 |

To get a deeper understanding of the type of information encoded within deep representations, we created an evaluation task of recovering the label of a given protein from the labels of its nearest neighbours for a cross-section of various ontologies. If the label is correctly recovered, it indicates that the representation is consistent within this ontology (Fig. 2). This paper focuses on investigating the representation and its features, not aiming at creating a universal label predictor.

**Figure 2.**
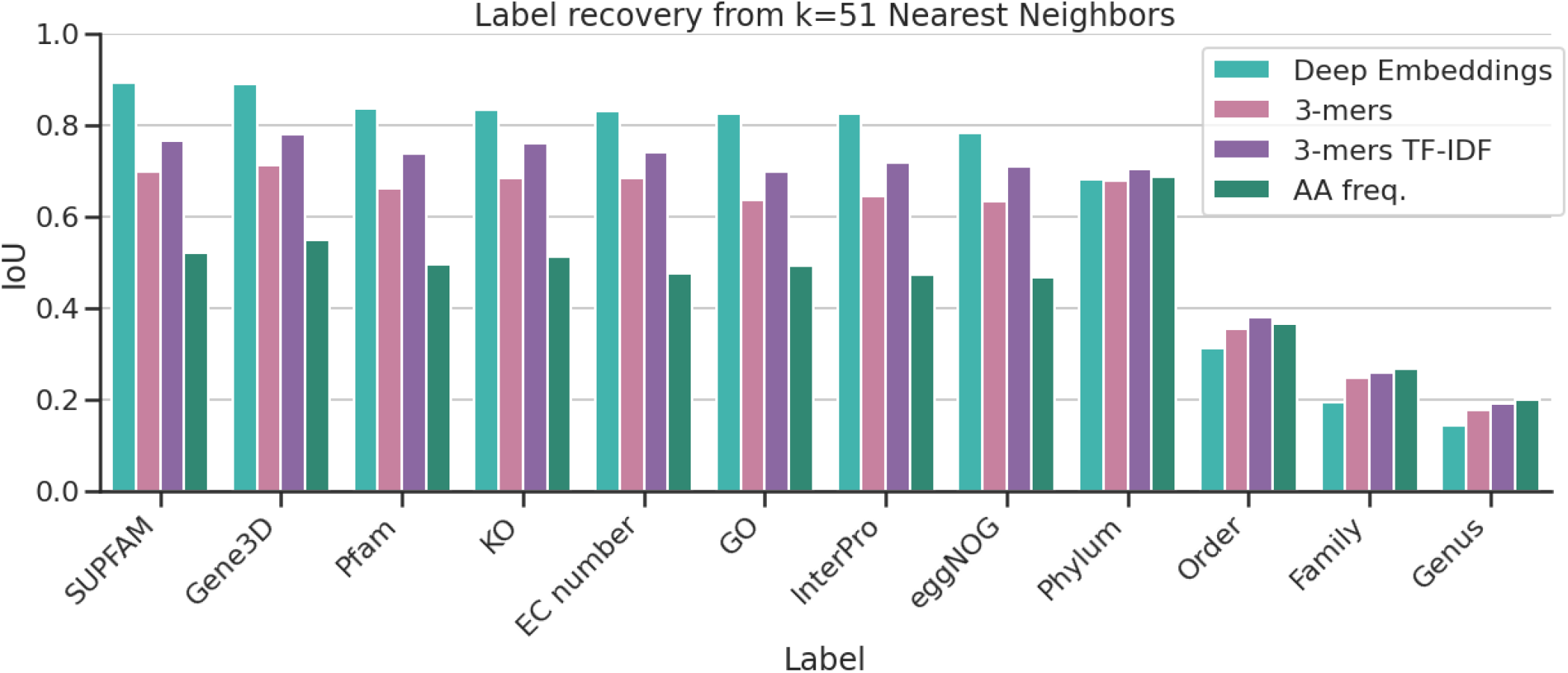
The degree of correctness in the recovery of labels using deep, k-mer-based or amino acid frequency representations. The measure of the recovery is Intersection over Union (IoU) between original labels and a set of labels from 51 nearest neighbors.

To evaluate the consistency of the representation, we selected a number of ontologies from Bacterial SwissProt, related to Function, Structure, or organism of Origin (Table 1). The ontologies significantly vary in the number of classes and Bacterial SwissProt coverage. Hence, the recovery task for each ontology may have a varying degree of difficulty. For this reason, we compared deep embeddings to general scalable sequence-based representations that do not use deep learning. Those baselines are 3-mers embeddings, 3-mers with term frequency-inverse document frequency (TFIDF) transformation, and amino-acid frequencies vecto (see Methods), similarly to seminal works in this field^35,38,43^.

In order to measure label recovery performance of our and baseline representations, we used a cross-validation-based approach. We removed labels of 20% randomly selected proteins in the dataset. Next, we trained a k-Nearest-Neighbor (kNN) classifier. Then, for every protein without a label, we predicted its label based on a majority vote from k nearest neighbors (k=51). We repeated this procedure 5 times.

Many proteins are annotated with more than one label within each ontology (for example, a protein may have multiple Pfam domains). To overcome this challenge, we used the Intersection over Union (IoU) metric. IoU is the ratio between the correctly predicted labels and the union of all predictions with all ground-truth labels for given protein (1). IoU ranges between 0 and 1, where 1 means perfect label recovery. For single-label tasks, IoU reduces to accuracy.

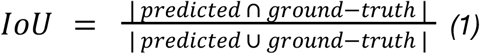

### Embedding performance on structure-, function- and taxonomy-related ontologies

Despite a varying number of classes in each task, the results from all ontologies unrelated to taxonomy were similar (Fig. 2). This suggests a comparable degree of difficulty between them, possibly due to the correlations amid labels (e.g. KOs are correlated with Pfam domains). The performance of all representations drops for taxonomic labels, esp. genus, family, and order (Fig. 2).

### Structure- and function-related ontologies

The deep representation generated performs best at recovering labels from ontologies based on protein structures (Gene3D, SUPFAM), while function- or domain-related ontologies obtained a slightly lower metric. These results indicate that the representation space primarily encodes the structure of proteins and secondarily the function of the protein, which is a structure derivative^47^.

The embedding model was trained to predict the best fitted amino acid in the protein sequence given all previous residues in the sequence. However, the results suggest that the model implicitly learnt to approximate the secondary and tertiary structure of the protein to improve in the training objective. It is reasonable as structural elements (eg. alpha-helix or beta-sheet) determine preferred, or the most suitable amino acids.

To this day, the most prevalent methods used for function annotation to proteins are based solely on sequence alignment. This approach is based on the hypothesis that proteins of similar sequences are usually homologous and thus, have a similar structure and function. But it is not always the case, hence the hypothesis is frequently misleading. On the other hand, embeddings, which are inherently structure-related, could enable more accurate function annotation tools.

### Taxonomy-related ontologies

In all representation spaces, predicting taxonomic origin is more difficult than functional characteristics with an exception for the highest taxonomic rank - phylum (Fig. 2). However, k-mers-based embeddings surpass deep ones in this task. K-mers are based solely on protein sequence, so they can be more biased towards the organism of origin, rather than the functional aspects of proteins^48,49^, (https://www.eb.tuebingen.mpg.de/protein-evolution/protein-classification/). However, in the process of evolution, genes undergo displacement, duplication, and horizontal transfer, which causes an increase in the taxonomic distance between protein sequence and its organism of origin. In the practice of taxonomy assignment genes need to be universal, conserved and not undergo frequent horizontal gene transfer ^50^. In general, protein sequence is a poor indicator of the organism of origin. We also see this considering both models’ performance on EggNOG ontology. It is combining information about function and taxonomy, achieving results between function- and taxonomy-related ontologies.

### Low dimensional representation of protein sequence space goes beyond sequence similarity

Representations learned by deep models are information-rich, but more difficult to understand due to the high dimensionality of the embedding. Further reduction in dimensionality with UMAP allows us to plot and visually interpret the embedding space built by the model.

### Deep embedding model creates a functionally structured representation space

To better understand which proteins were the easiest to recover based on the embedding, we defined Recovery Error Rate as *1 - average IoU* metric obtained on each protein across all ontologies. The use of this metric enabled us to localize regions with low & high Recovery Error Rates (Fig. 3). In Fig. 3A, we show that proteins with low Recovery Error Rates are located in smaller clusters, while proteins with high error rates are concentrated in the center of the UMAP visualization.

**Figure 3.**
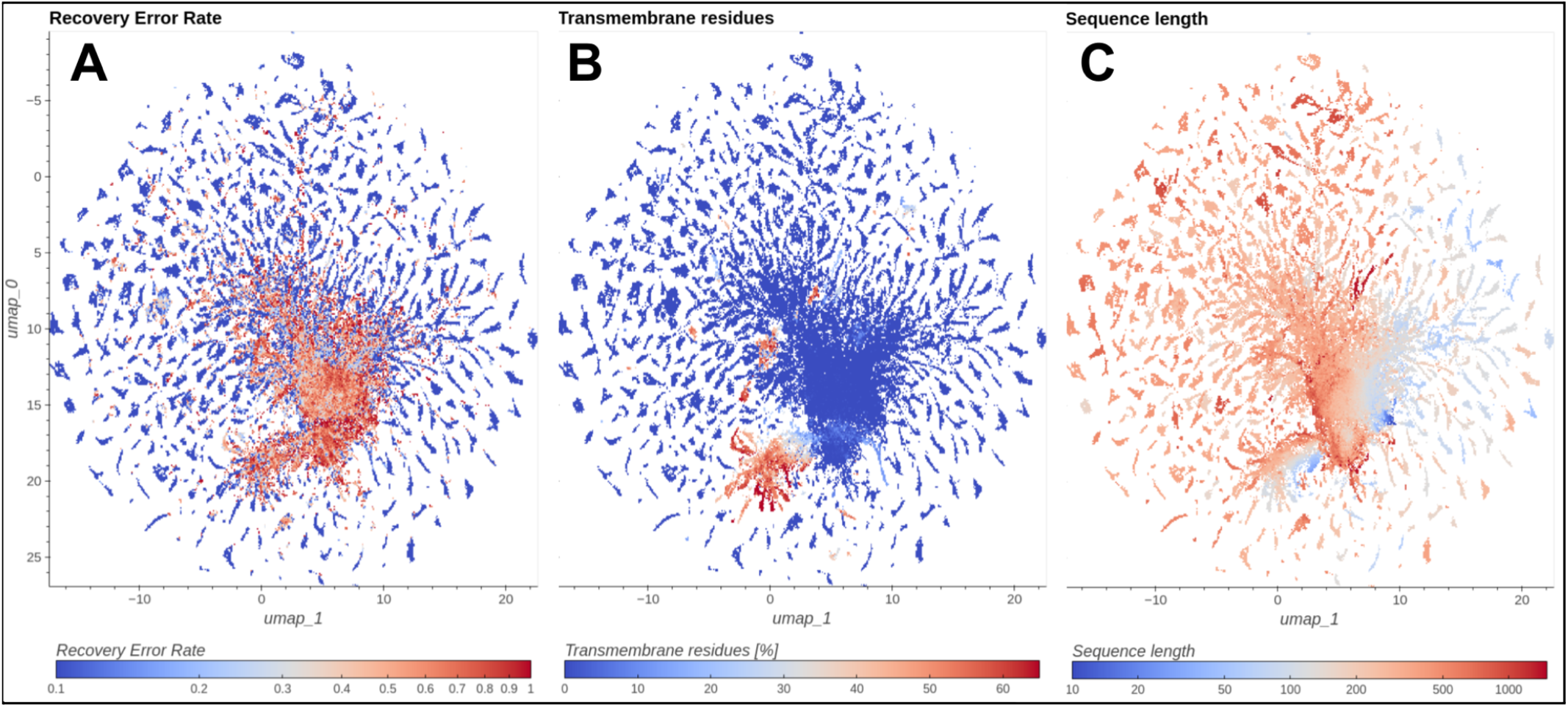
UMAP visualization of Bacterial SwissProt embeddings. (**A**) Proteins colored by Recovery Error Rate. (**B**) Proteins colored by percentage of transmembrane residues. (**C**) Proteins colored by sequence length.

To investigate the functional structure of the representation space, we overlay it with labels defined by Kegg Orthology ID (KO) (Fig. 4). The proteins that do not have a KO assigned are colored in grey - we see that they are placed in the central part of the plot. Most of the proteins are clearly clustered by their functional annotation. Furthermore, by focusing on specific space locations, we can see that close KO clusters share other functional features: domains (Fig 4A & 4B), EC number class (Fig 4A & 4D), or structural and molecular features (Fig 4E). It suggests that the deep representation does not focus only on one functional ontology, but rather on an abstract protein function defined on many levels. The visualisation explains the high label recovery results and expands analogous analysis conducted on a smaller scale with only 25 COGs^38^. Compared to the k-mer based representation, the deep representaion is significantly more structured (Supplementary Fig. 1).

**Figure 4.**
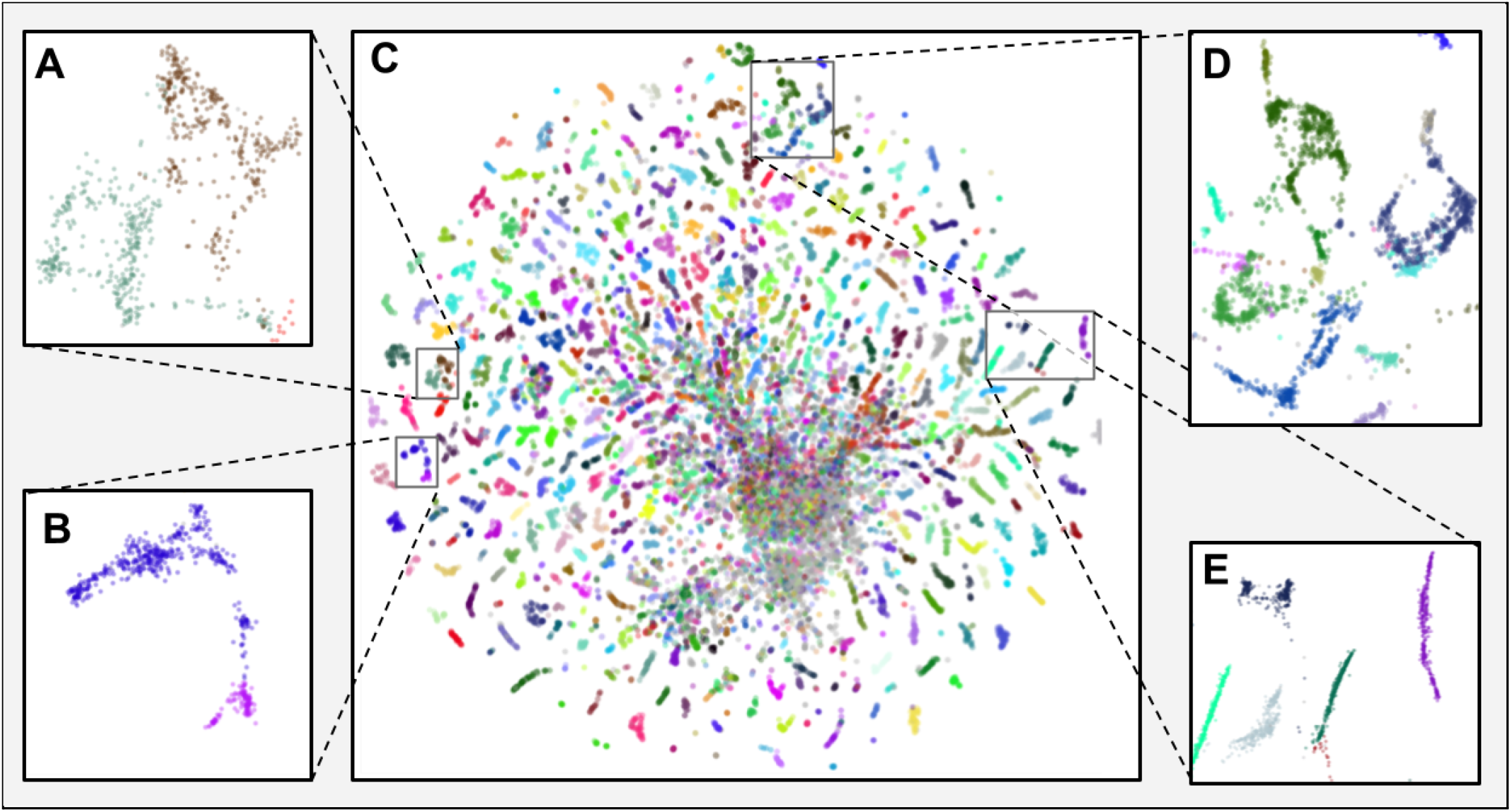
Deep embeddings UMAP projection of Bacterial SwissProt colored by KO. (A) transferase proteins that share the same Pfam domain and belong to the EC 2.5.1 class - UDP-N-acetylglucosamine 1-carboxyvinyltransferase (K00790) in dark green, 3-phosphoshikimate 1-carboxyvinyltransferase (K00800) in brown (B) GTP binding proteins sharing Pfam domains - Elongation Factor G (K02355) in purple, Peptide chain release factor (K02837) in pink. (C) all Bacterial SwissProt proteins (D) proteins that belong to the tRNA ligases class (EC 6.1.1) - Cysteine (K01883), Arginine (K01887), Glutamate (K01885), Glutamine (K01886), Glycine (K01880), Valine (K01873), and Isoleucine (K01870) (E) ribosomal proteins - 30S ribosomal protein S1 (K02961) in light green, 50S ribosomal protein L14 (K02874) in light blue, 50S ribosomal protein L36 (K02919) in black, 50S ribosomal protein L35 (K02916) in dark green, and 50S ribosomal protein L15 (K02876) in purple. For a detailed description of the protein set see Supplementary Table 2. The whole space can be interactively explored in our application (https://Drotein-exDlorer.ardigen.com).

We hypothesize that the regions of high Recovery Error Rate are occupied by rare proteins. Rare proteins form small functional classes in Bacterial SwissProt, and the smaller the functional class is, the more difficult it is to predict the label based on its neighbors. Additionally, their potential insufficient representation in the training set makes it difficult to model their sequences, as the embedding model can learn certain patterns only if they are shared by a sufficient number of proteins in the training dataset. Indeed, we observed that the Recovery Error Rate is negatively correlated (r=-0.715, N=200,115) with the log-average size of the functional class the protein belongs to (See Supplementary Fig. 2). Moreover, we noticed an increased frequency of the occurrence of words: *‘Uncharacterized’*, *‘Putative’* and *‘Probable’* in SwissProt descriptions of error-causing proteins (25% for Recovery Error Rate = 1 vs. 4% for Recovery Error Rate = 0, See Supplementary Fig. 3), indicating less characterized proteins.

### Short and transmembrane proteins

The embedding model is sensitive to the length of the protein (Fig. 3C) and a significant number of short proteins is present in the central, lesser understood part of UMAP visualization. Short proteins (≤50 residues), underestimated for a long time, gained interest in recent years when it was discovered that they are involved in important biological processes such as cell signaling, metabolism, and growth^51^. The presence of a high Recovery Error Rate region might be a result of insufficient information on small proteins, which are still underrepresented in databases. Following Sberro et al., based on the NCBI GenPept database, over 90% of small protein families have no known domain and almost half are not present in reference genomes^52^.

Transmembrane proteins constitute approx. 30% of all known proteins. Unlike globular proteins, they are on average larger and must exhibit a pattern of hydrophobic residues to fit into the cell membrane^53^. In order to define transmembrane proteins we used a transmembrane score (a percentage of transmembrane residues) adopted from Perdigão et al.^54^. In Figure 3B, we can see that the model separates transmembrane proteins well, which is in line with previous research on deep protein representations^42,55^. However, part of transmembrane proteins lie within the high-recovery error region of the UMAP plot. Despite substantial pharmacological and biological relevance, they are less understood and underrepresented in databases, as structural experiments on them are difficult to conduct.

Lower ECE loss obtained on metagenomic proteins (compare *Alignment-free deep protein embeddings represent structure- and function-related ontologies* section) suggest that the deep embedding model trained on a more general catalog of metagenomic proteins (UHGP) is less biased towards well-known model organisms, hence, better suited for rare, short or transmembrane proteins.

### A sample use case - phosphotransferases (EC 2.7.2)

To demonstrate the use of embedding representation in a real-life scenario, we used a group of phosphotransferases. We have chosen them due to their importance in maintaining the human gut microbiome homeostasis. Acetate, butyrate, and propionate kinases are especially crucial in the process of forming short-chain fatty acids (SCFAs). SCFAs are produced in the colon by bacteria during the fermentation of resistant starch and non-digestible fibers. Their lowered level is often observed in patients suffering from irritable bowel diseases (IBD) such as Crohn’s disease and ulcerative colitis. SCFAs serve as an important fuel for intestinal epithelial cells and participate in preserving gut barrier integrity. Recent findings indicate their role in energy metabolism (lipid metabolism), immunomodulation, regulation of intestinal epithelial cells, proliferation and cancer protection. Although promising, the research has been conducted mainly on murine or *in vitro* models, thus the results have to be interpreted with caution^56–58^.

Proteins classified as phosphotransferases were chosen based on their EC number. We decided to use this annotation as EC numbers are a manually assigned nomenclature that describes enzymes based on the chemical reactions they catalyse. Their hierarchical structure allows for a fine-grained analysis. Proteins described by EC 2.7.2 class represent phosphotransferases with a carboxyl group as an acceptor. We used eight EC 2.7.2 subclasses available at Bacterial SwissProt: EC 2.7.2.1 (acetate kinase), EC 2.7.2.2 (carbamate kinase), EC 2.7.2.3 (phosphoglycerate kinase), EC 2.7.2.4 (aspartate kinase), EC 2.7.2.7 (butyrate kinase), EC 2.7.2.8 (acetylglutamate kinase), EC 2.7.2.11 (glutamate 5-kinase) and EC 2.7.2.15 (propionate kinase).

We examined the domain architecture of EC 2.7.2 proteins using the Pfam database. The domain architecture is the main structure that defines a protein’s function. We found that four domain architectures were dominant among analysed proteins. 31% of analyzed proteins contained one amino acid kinase domain (PF00696), 29% of proteins had one phosphoglycerate kinase domain (PF00162), 20% contained one acetate kinase domain (PF00871), and 18% of proteins had two coincident domains PF00696 & PF01472, i.e., amino acid kinase domain and PUA domain.

In total, we study 1,302 proteins exhibiting eight unique specific functions (ECs) and four distinct domain architectures (See Supplementary Table 3). Different domain architectures suggest that these proteins have different amino acid sequences and would be difficult to identify as similar with baseline bioinformatic methods based on sequence similarity alone.

To investigate how accurately the embedding representation reflects the functional relationships between the proteins, we visualized them using UMAP (Fig. 5A). Almost all proteins were grouped according to their domain architecture, and proteins with similar domain architectures, such as proteins having only PF00696 domain and proteins having two domains PF00696 & PF01472, were also placed closer to each other. Despite clear domain-based grouping, proteins that share the same domain architecture, but catalyze different chemical reactions, are separated. The only exceptions are EC 2.7.2.1 and 2.7.2.15. One possible explanation for this exception is that these two enzymes can share substrates for their activity. Acetate kinases (EC 2.7.2.1) can accept propionate as an alternative substrate, and propionate kinases (EC 2.7.2.15) can accept acetate. Moreover, both EC 2.7.2.15 and EC 2.7.2.1 play essential roles in the production of propionate in bacteria^59^. The only inconsistency we can note are two butyrate kinase proteins (Figure 5A cyan circles; PF00871) that were placed far from their counterparts.

**Figure 5.**
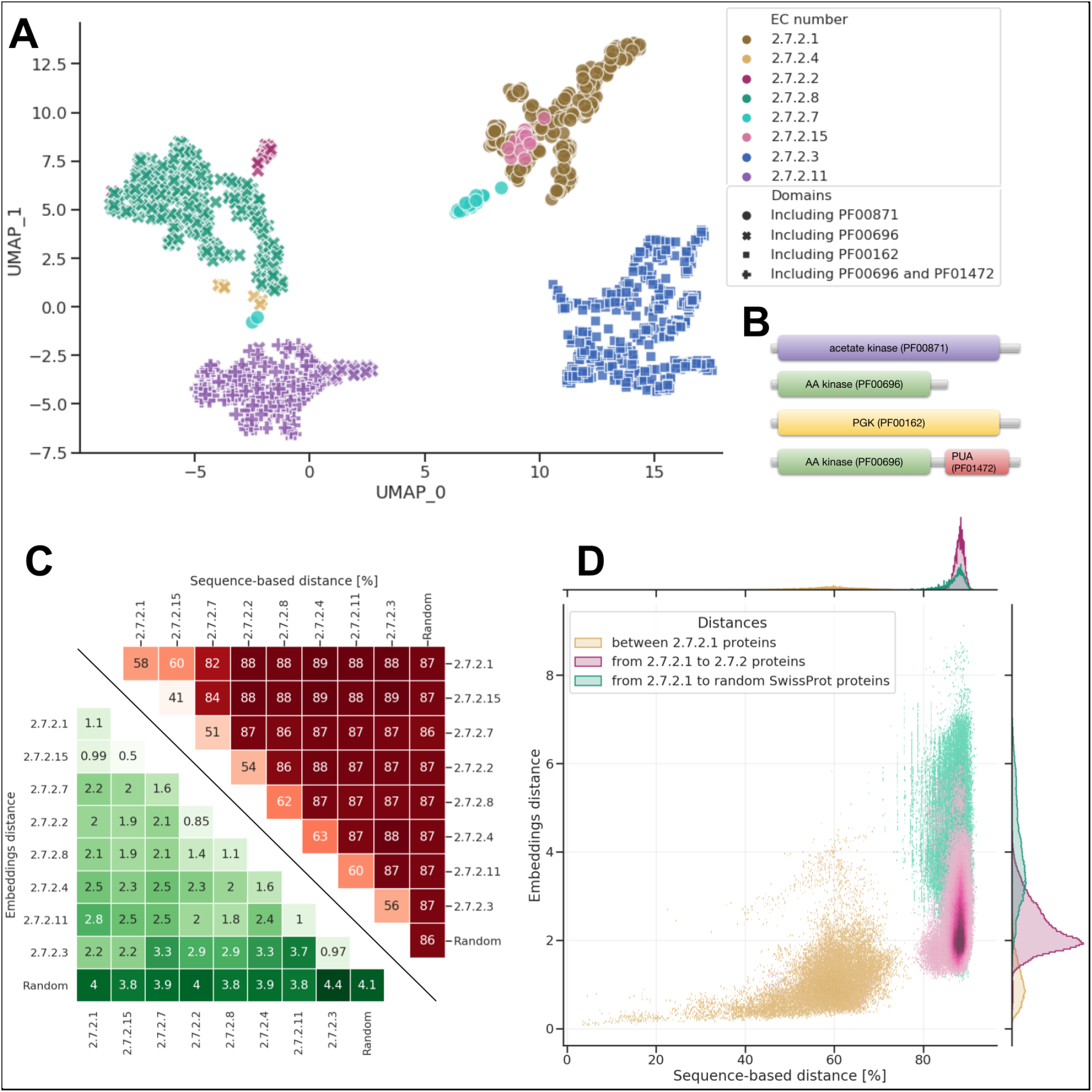
Visualization of EC 2.7.2 proteins in the deep embedding space. (**A**) Deep embeddings of EC 2.7.2 proteins visualized with UMAP. (**B**) Domain architecture of EC 2.7.2 (**C**) The mean distance between EC 2.7.2 proteins and 500 random proteins from the SwissProt space with distinction between embedding-based distance (green) and ClustalO distances (red). Values for both methods were calculated as averages of pairwise distances between all proteins within given clusters. The mean embedding-based distance between EC 2.7.2 proteins is significantly smaller compared to the distance between EC 2.7.2 proteins and 500 random proteins. Not only proteins from the same cluster group are closer to each other but also proteins from different EC 2.7.2 clusters are located significantly closer to proteins from other EC 2.7.2 clusters than to random proteins. Mean distance between proteins calculated using ClustalO does not reflect the clear separation between EC 2.7.2 and random proteins. Mean ClustalO distance between proteins from the same cluster is smaller than between EC 2.7.2 and random proteins, however ClustalO does not bring proteins closer from different EC 2.7.2 clusters. (**D**) Comparison of embedding-based and sequence-based distance (ClustalO) to EC proteins 2.7.2.1. The distances were divided into those within the protein group EC 2.7.2.1, from EC 2.7.2.1 to other EC 2.7.2 proteins, and from EC 2.7.2.1 to randomly selected proteins. The embedding-based, as opposed to the sequence-based distance, differentiates the distances from EC 2.7.2.1 to other members of EC 2.7.2 and from EC 2.7.2.1 to random proteins.

To further analyze two outlying butyrate kinases, we inspected sequences of outlier’s domains sequence and compared it to sequences of PF00696 and PF00871 domains. We performed multiple sequence alignment (MSA) of outlying protein’s domains, the PF00696 domains and the PF00871 domains sequences (see Methods). MSA results showed that outlying protein’s domain sequences are distant from PF00871 domain sequences from further proteins, including other butyrate kinases. However, outliers were more closely aligned with sequences that belonged to PF00696 domains (Supplementary Data 1-3). We hypothesize that the domain sequence of the two outlying proteins is more similar to PF00696 domain than PF00871, which made UMAP place them closer to the former (Figure 5A).

We hypothesize that the deep representation reflects the functional similarities between proteins that are based on domain architecture (Pfam domains) or enzymatic activity (EC number). This emphasizes the significant advantage of deep embeddings, as they do not only focus on single, human-created ontology, such as e.g., EC numbers, but rather fuse all information to characterize proteins on multiple levels. It combines the strengths of approaches that focus on motifs, domains (Pfam), and 3D structure (GENE 3D) to understand protein function space comprehensively.

To better understand the differences between sequence-based distance and deep embeddings, we compared the Euclidean distance between EC 2.7.2 proteins and randomly chosen 500 proteins from the Bacterial SwissProt dataset. As a baseline, we selected sequence-based distance calculated with Clustal Omega^60^. The distance measure used by Clustal Omega for pairwise distances of unaligned sequences is the k-tuple measure.

The embedding-based distances within and between the EC 2.7.2 subclasses are smaller than to randomly selected proteins, which do not hold for the sequence-based distance (Fig. 5B & 5C). This proves that the embedding model can go beyond sequence similarity and find relations between proteins with significantly different sequences and domain architectures. This property enables searching for proteins that are similar on a more abstract level and in the future may improve the annotation coverage of microbial proteins.

## Discussion

The human microbiome plays a crucial role in human health, and changes in its composition can be related to various diseases, such as diabetes, cancer, or psychiatric disorders. To fully understand the complex relation between the microbiome and human health, it is necessary to look not just at the taxonomic level but also at a functional level. Despite various approaches to retrieve protein functions^61,62^, a large portion of microbial proteins remain functionally uncharacterized. This paper presents a novel approach of using the Bidirectional LSTM model to visualize and contextualize the microbial protein space. We show that our model accurately represents protein features related to structure and function, overcoming some limitations of standard bioinformatics methods such as HMMER or BLAST.

The deep learning model creates an abstract, numerical representation of proteins in an embedding space. This embedding encodes information from various protein ontologies and combines knowledge on protein structure and function, overcoming the limitations of methods based on sequence similarity. At the same time, generating the embedding for a given protein is much more efficient computationally than using bioinformatic methods. On top of that, the embedding is also more suitable for a large range of further downstream algorithms, such as classification, clustering and visualization. Combining embeddings with a dimensionality reduction method, such as UMAP, may enable creating a reference protein map and facilitate protein research.

One of the significant challenges that any data-driven solution must face is data bias. Our results indicate that using a catalog of metagenomic proteins (UHGP) for training made the model less biased towards well-known model organisms. Despite this, model validation required the use of experimentally verified data, which limited the scope of our validation to well-known proteins and prevented genuine validation on small or transmembrane proteins. We assume that with the growing interest in these proteins, their presence in the databases and number of their annotations will increase, which will allow for a more thorough validation.

We are witnessing rapid progress in both the deep learning field and in metagenomics, which generate massive amounts of data. We believe embedding models are an attractive alternative to database-bound, computationally intensive methods unsuitable for such influx of data. An appealing approach would be to join the strengths of computationally-cheap embedding models with other computational technologies that can accurately predict the features of individual genes (for example: protein 3D structure using AlphaFold^19–21^) and finally perform experimental validation on most promising targets. Such approach enables such efficient contextualization of metagenomic data and may be used to better understand the microbiome for health.

## Methods

### Embedding model training

In the training, we took advantage of the Unified Human Gastrointestinal Protein catalog clustered at 95% sequence identity (UHGP-95) to limit the impact of the most common sequences. Further clustering may improve the model^38^. UHGP-95 contains exactly 19,228,304 protein sequences, from which we randomly selected 5% to track training progress (validation set) and set aside another 5% for the final model evaluation (test set). The rest of the data (18,266,888 sequences) was used to train the model. All proteins were clipped to 1,500 amino acids.

We used a 3-layered Bidirectional LSTMs (BiLSTM) model with 1024 hidden units in each layer. We have chosen the LSTM architecture as it gave the best results in Remote Homology detection in the TAPE benchmark^37^ and achieved superior performance over Transformer-based architecture in the ProtTrans benchmark^42^. On the other hand, the most recent findings show the superiority of Transformer-based architectures^21,38,63^ in protein informatics. We assume that those and even further advances in deep learning, especially applied to protein sequences, will only improve results presented in our work.

The model was trained by the AdamW optimizer for 225,331 weight updates with a mini-batch of size 1024, which corresponds to 12 epochs and approximately 48 hours on 4 Tesla V100 GPUs. The learning rate was set to 1e-3, except the first 8,000 steps that were used as a warmup. The process was implemented in the PyTorch library, based on the TAPE benchmark^37^ repository (https://github.com/songlab-cal/tape).

### Computing embeddings

To obtain a vector representation of a protein (embedding) from the BiLSTM model, we extracted vectors of hidden states for each amino acid and averaged them. This is in contrast to natural language processing practice, which uses the hidden state vector corresponding to the last word (here it would be the last amino acid) rather than the average representation of all words. However, there is evidence suggesting the superiority of averaged presentation in the field of protein processing^43^. This may be due to the fact that proteins are usually much longer than sentences, and LSTM-based models cannot fit the whole amino acid sequence in just one state.

### Validation on metagenomic proteins

The UHGP catalog contains data publicly available as of March 2019, thus for the validation we have selected ten samples (See Supplementary Table 1) from a dataset published in May 2020 (PRJEB37249)^64^. We assembled samples using MEGAHIT v1.2.9^65^ and retrieved protein sequences using prokka v1.14.6^66^ on obtained contigs. Samples contained between 22640 and 108120 protein sequences, 50035 on average. We measured ECE loss on each sample separately and average values to obtain final results. For comparison we trained a model on Pfam database v32^67^ - the same as used in TAPE benchmark^37^.

### Bacterial SwissProt

For evaluating the properties of the embedding space, we used the UniProtKB/Swiss-Prot 2019_02 database with 562,438 protein entries. For every entry, we parsed taxonomy lineage and functional labels (Table 1). Only proteins from the Bacteria domain were selected, leaving 331,523 proteins.

To remove near-identical protein sequences, we deduplicated the remaining set using *mmseq2 easyclust* with an identity threshold set to 97% and coverage set to 0.8. Removing duplicates ensured no cliques in the kNN graph, which we used in the kNN label recovery and UMAP visualizations. Cliques would lead to trivial solutions during kNN classification and “lonely islands’’ in UMAP visualizations.

After the deduplication step, we obtained 201,622 proteins, and this set we named Bacterial SwissProt.

### Baseline representations

For a general sequence-based baseline representation, we used the bag of k-mers method^68^, which produces embedding for a protein by the following procedure: (a) generate all possible k-mers (subsequences of length k) from protein sequence, (b) count occurrences of each possible k-mers in the sequence, (c) sort counts alphabetically by k-mers sequence. Sorted counts form a vector representing the sequence.

Higher k leads to more specific representation but increases dimensionality, which is equal to the number of all possible k-mers (N=20^k^). In our work, we choose k=3, which resulted in 8,000-dimensional vectors.

We also included a variant of 3-mer representation with term frequency-inverse document frequency (TFIDF) transformation, which accentuate rare k-mers. To complement k-mer-based baselines we added a representation of the amino acid frequencies vector.

### Label recovery

For the analysis, we used the Bacterial SwissProt described above. We generated deep and k-mer representations for each protein. Next, we reduced the dimensionality of both representations to 50 using the Principal Component Analysis (PCA) algorithm (Fig. 1B).

We narrowed down the set of analyzed proteins to only those with assigned labels in given ontology for each ontology analyzed. We divided these sets of proteins into five equal parts to estimate recovery efficiency through 5-fold cross-validation. For every fold, we constructed a kNN graph (https://github.com/lmcinnes/pynndescent) of the data from the four remaining folds. The graph was then used to predict classes for each protein in the fold, by querying the nearest proteins (N=51) and propagating their labels as a prediction. As the protein can be assigned to many classes (multi-label classification), we used the Intersection over Union (IoU) metric. A higher number of neighbors (N) taken into account causes the results to extend beyond the immediate neighborhood, which usually contains highly similar proteins. At the same time, the larger N is, the more challenging it is to predict a label for small classes, and it even becomes impossible for the classes smaller than N / 2 proteins. We choose 51 to balance these two properties.

### UMAP visualizations

To visualise protein embedding space, we further reduced dimensionality of the PCA Reduced Embeddings with UMAP (Uniform Manifold Approximation and Projection; https://github.com/lmcinnes/umap), a nonlinear dimensionality reduction method.

UMAP was chosen over another common nonlinear dimensionality reduction method, t-SNE (t-distributed Stochastic Neighbor Embedding), as it preserves more of the global structure with superior run time performance^69^.

### 2.7.2 cluster analysis

#### Selecting proteins

Proteins assigned to EC 2.7.2 subclass were chosen for the analysis. In the analysis, we used 8 available EC 2.7.2 sub-subclasses out of 14, as our bacterial dataset lacked proteins described by 6 other sub-subclasses. Sub-subclasses used in this analysis are EC 2.7.2.1 (acetate kinase), EC 2.7.2.2 (carbamate kinase), EC 2.7.2.3 (phosphoglycerate kinase), EC 2.7.2.4 (aspartate kinase), EC 2.7.2.7 (butyrate kinase), EC 2.7.2.8 (acetylglutamate kinase), EC 2.7.2.11 (glutamate 5-kinase) and EC 2.7.2.15 (propionate kinase). We assigned a Pfam ID to each protein using mapping available in SwissProt. 4 domain architectures were found dominant among 1,302 analysed proteins. 31% of analyzed proteins contained one amino acid kinase domain (PF00696), 29% had one phosphoglycerate kinase domain (PF00162), 20% one acetate kinase domain (PF00871) and 18% had two coincident domains (PF00696 and PF01472), i.e. amino acid kinase domain and PUA domain.

We visualized EC 2.7.2 proteins in the same manner as described above in *UMAP visualizations*.

#### Comparison to sequence (Clustal Omega for distance matrix)

To infer about the ability of the embedding model to group more closely proteins sharing a function, we compared the distance between EC 2.7.2 proteins and 1,000 randomly chosen proteins from the Bacterial SwissProt database. We wanted to analyze if embeddings distance between proteins is compatible with corresponding amino acid sequence distance. embedding distance was calculated as an Euclidean distance between 50 PCA components. Those 50 PCA components are the result of dimensionality reduction of 2,048 protein embeddings, generated by the model. Sequence distance was calculated using Clustal Omega ^70^, a bioinformatic tool for multiple sequence alignment. This tool takes a fasta file with unaligned protein amino acid sequences as input and calculates percent of sequence identity between those sequences giving a pairwise distance matrix. The distance measure used by Clustal Omega for pairwise distances of unaligned sequences is the k-tuple measure.

#### Outliers analysis

To infer sequence similarity we performed multiple sequence alignment (MSE) between outlying protein’s, PF00696 and PF00871 domain sequences. HMMER software^62^ was used to find domain positions in each protein. Biopython^71^ was used to extract domain from protein sequence. MSE was performed using Clustal Omega^60^. We performed three MSE: a) outlying butyrate kinases vs other butyrate kinase proteins, b)outlying butyrate kinases vs PF00696 domain sequences from proteins containing that domain, c) outlying butyrate kinases vs PF00871 domain sequences from proteins containing that domain. We visualized the alignment using Jalview^72^.

## Acknowledgements

We would like to thank: Monika Majchrzak-Górecka and Paweł Biernat for reviewing the draft of the manuscript and for the helpful discussion; Karol Horosin and Mateusz Siedlarz for technical help with the interactive visualization; Michał Kowalski for technical help with GPU servers.

## Author contributions statement

**KO** - conceived the study, conducted analyses, analysed the data, wrote the paper

**ZK** - analysed the data, wrote the paper

**JM** - discussed the results

**AB** - helped supervise the project

**KMZ** - discussed the results, helped supervised the project

**TK** - conceived the study, wrote the paper, helped supervise the project

All authors discussed the results and contributed to the final manuscript.

## Code availability

Code used in the analyses is available at https://github.com/ardigen/microbiome-protein-embeddings

## Supplementary information

**Supplementary Table 1.** Summary of samples selected from PRJEB37249 project for metagenomic validation of the deep embedding model.

| SampleId<br>(Run Accession) | n. of proteins | ECE |  |
| --- | --- | --- | --- |
|  |  | UHGP model | Pfam model |
| ERR4086424 | 38,410 | 11.421 | 15.4 |
| ERR4086425 | 55,508 | 10.932 | 15.173 |
| ERR4086426 | 41,077 | 10.842 | 15.176 |
| ERR4086427 | 51,965 | 10.813 | 15.131 |
| ERR4086428 | 61,933 | 10.81 | 15.123 |
| ERR4086429 | 59,977 | 10.779 | 15.103 |
| ERR4086430 | 108,120 | 10.204 | 17.066 |
| ERR4086431 | 23,021 | 11.326 | 15.168 |
| ERR4086432 | 22,640 | 10.575 | 14.86 |
| ERR4086433 | 37,703 | 11.528 | 15.201 |
| mean | 50,035 | 10.923 | 15.340 |
| std | 24,711 | 0.40 | 0.62 |

**Supplementary Table 2.**
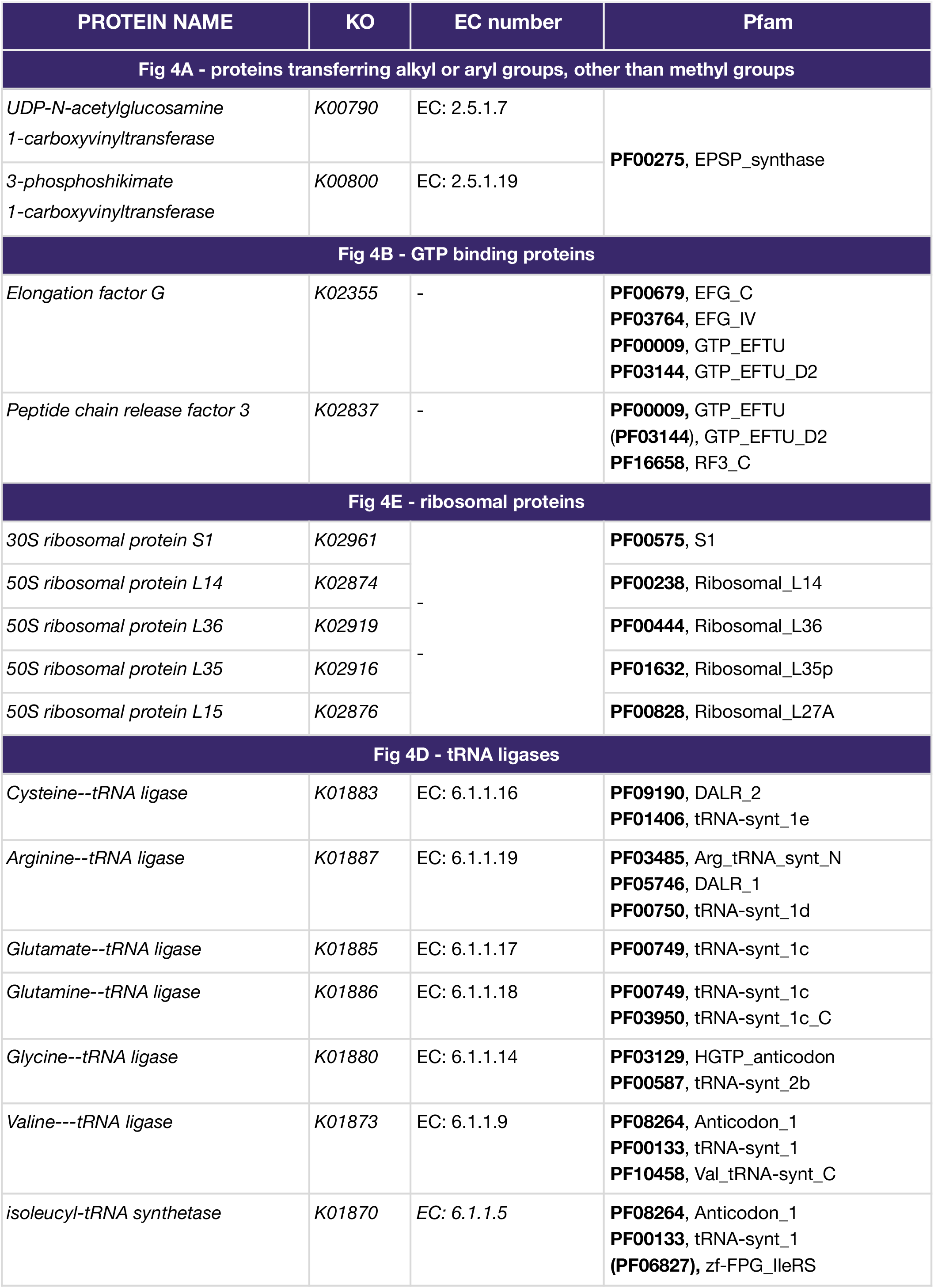
Annotation of proteins that are clustered in UMAP visualization to functional databases such as KEGG, GO, Pfam and annotation to EC number. We can see that proteins within a cluster share annotations.

**Supplementary Table 3.**
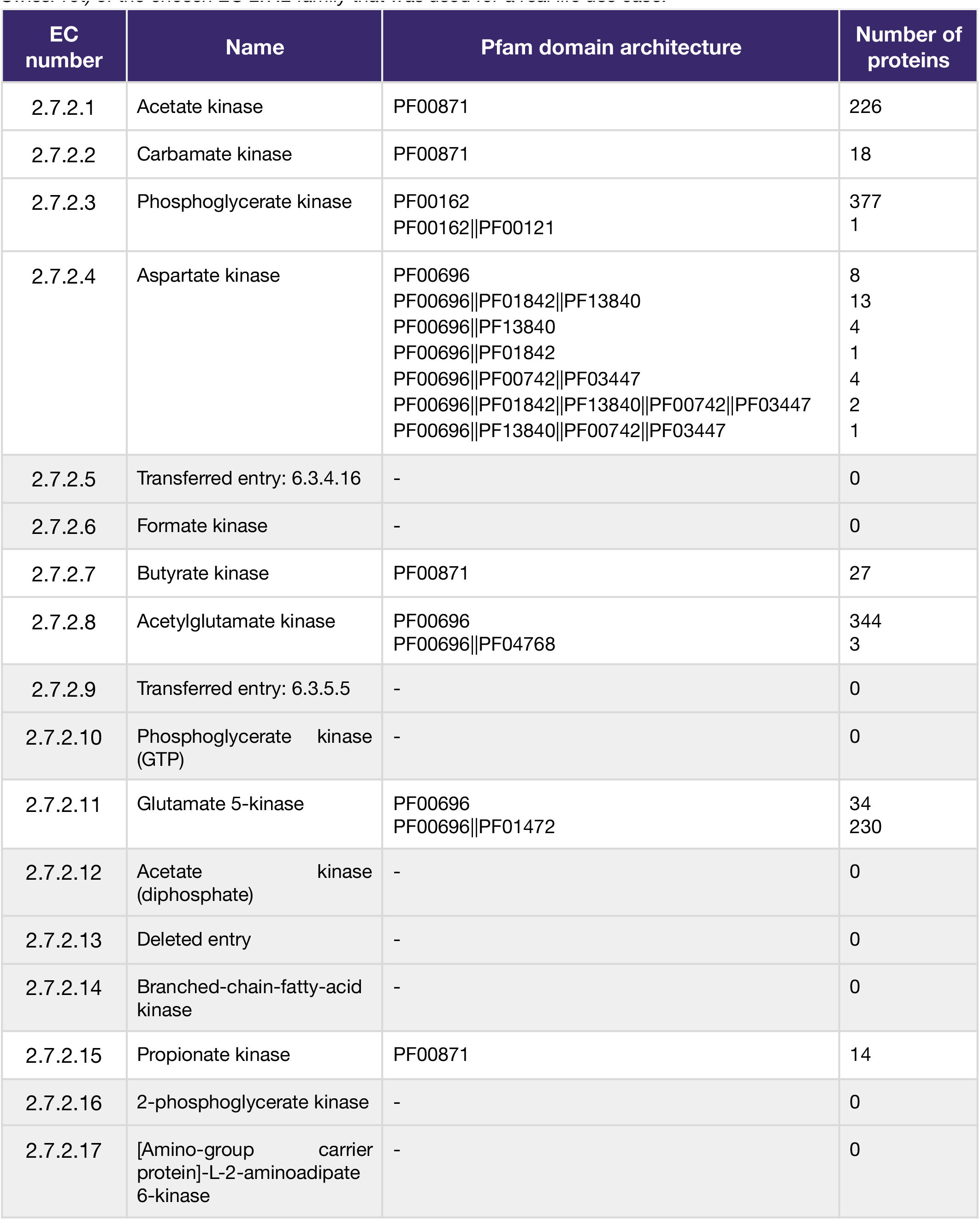
Description (i.e.: EC number, name, domain and number of proteins in Bacterial SwissProt) of the chosen EC 2.7.2 family that was used for a real life use case.

**Supplementary Figure 1.**
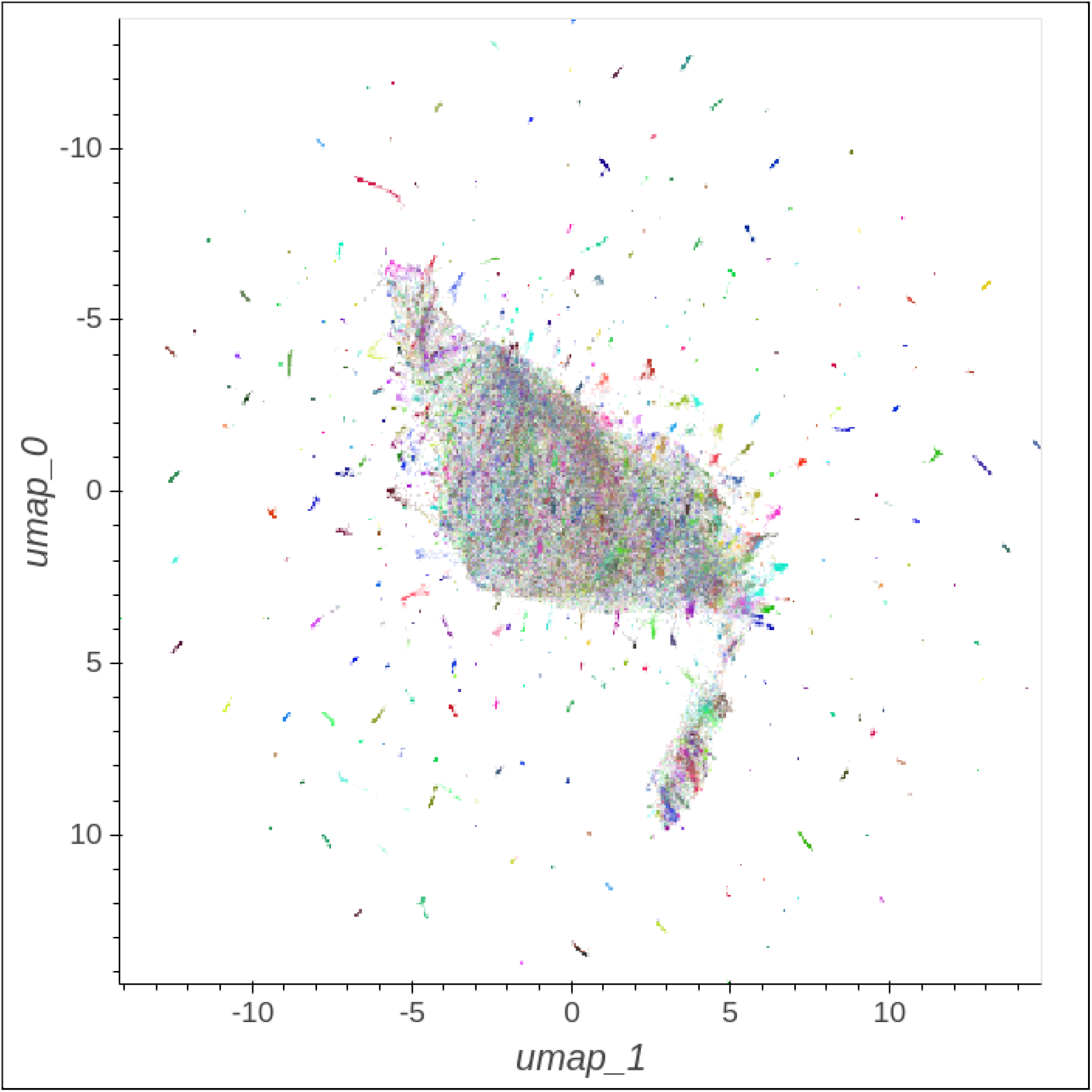
UMAP visualization of the k-mer protein representations space colored according to Kegg Orthology ID (KO). We see separate groups of proteins, however, most of them are mixed up in the middle.

**Supplementary Figure 2.**
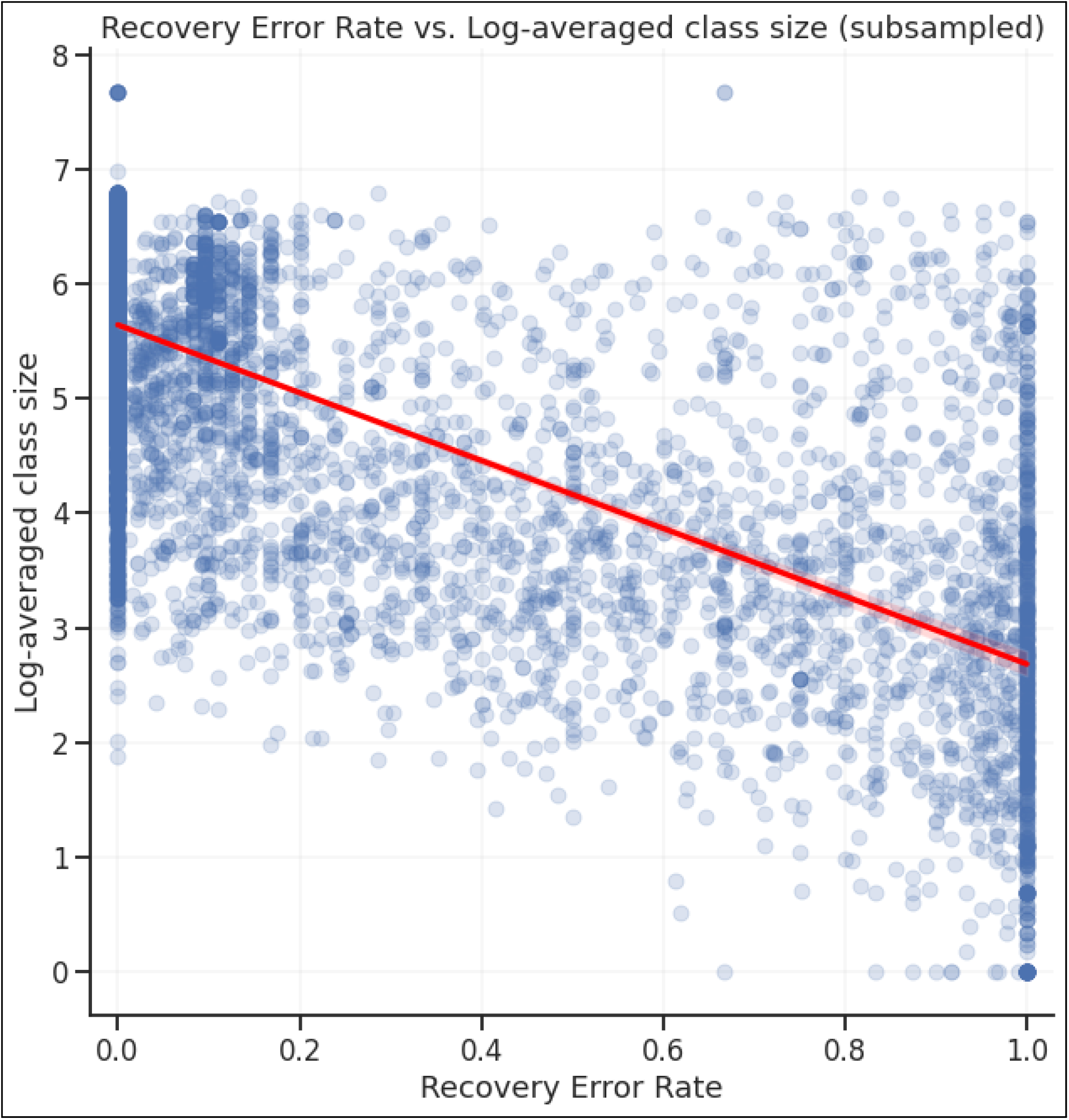
Relationship between Recovery Error Rate and the size of the class to which the protein belongs.

**Supplementary Figure 3.**
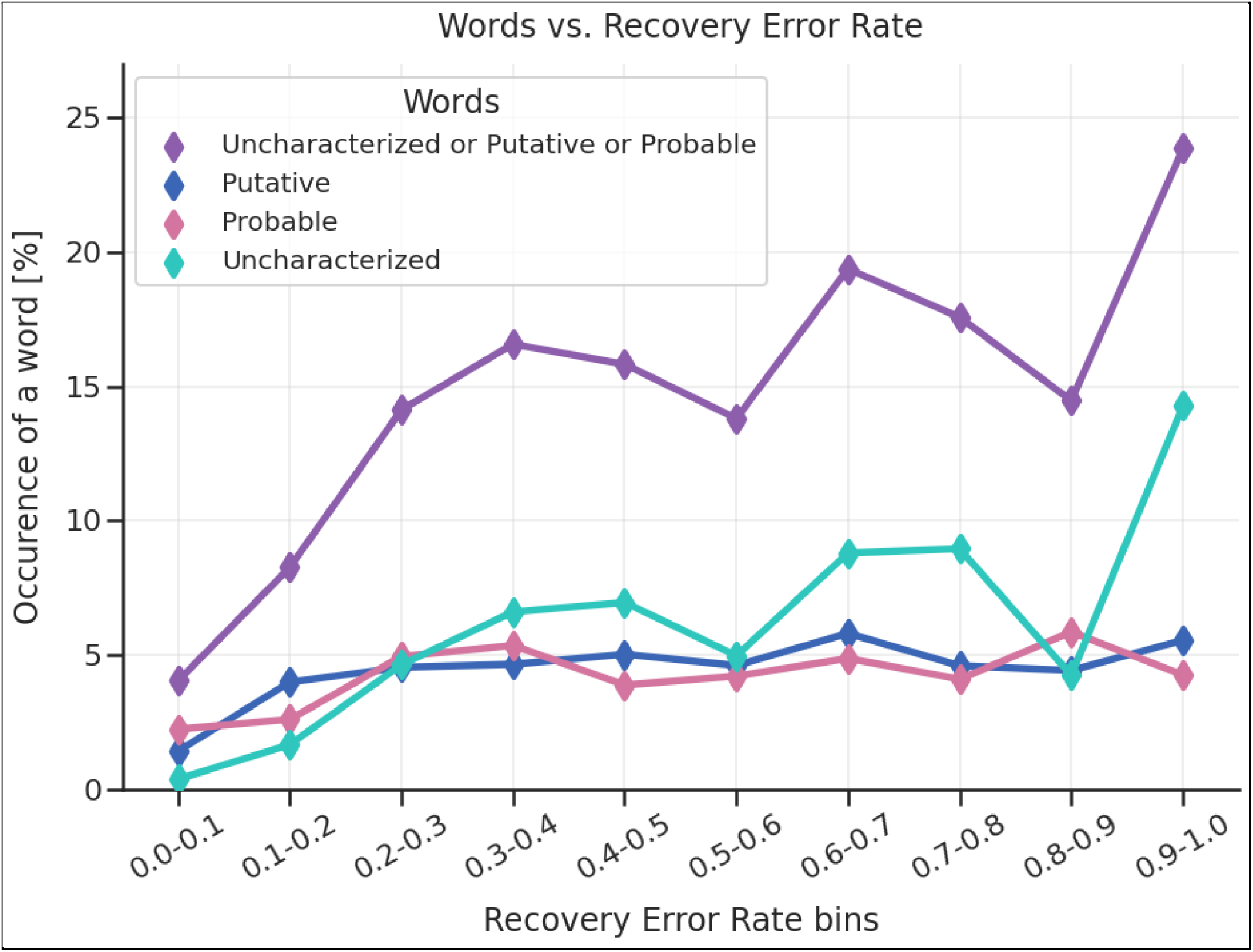
The relationship between Recovery Error Rate and the occurrence of the words “Uncharacterized”, “Putative”, or “Probable”.

**Supplementary Table 3.**
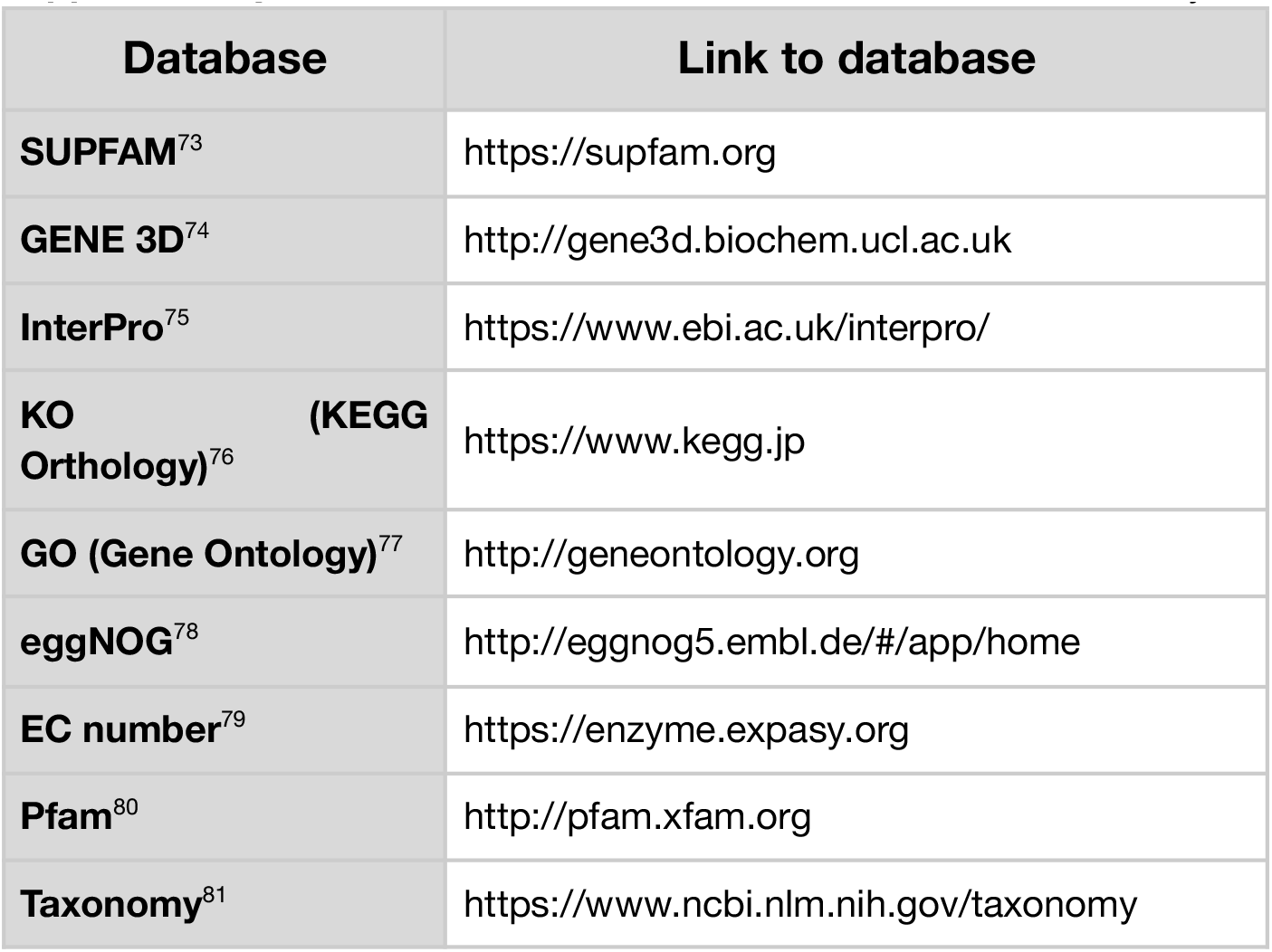
References to the databases used in our analyses.

