## Supplementary Data 2 for "Deep embeddings to comprehend and visualize microbiome protein space"

**Supplementary data 2.** Domain sequence alignment between two outlying proteins (BUK_OCEIH and BUK_DEIRA) and PF00696 domain sequences from other phosphotransferases (EC 2.7.2.8, 2.7.2.2, 2.7.2.4 and 2.7.2.11)

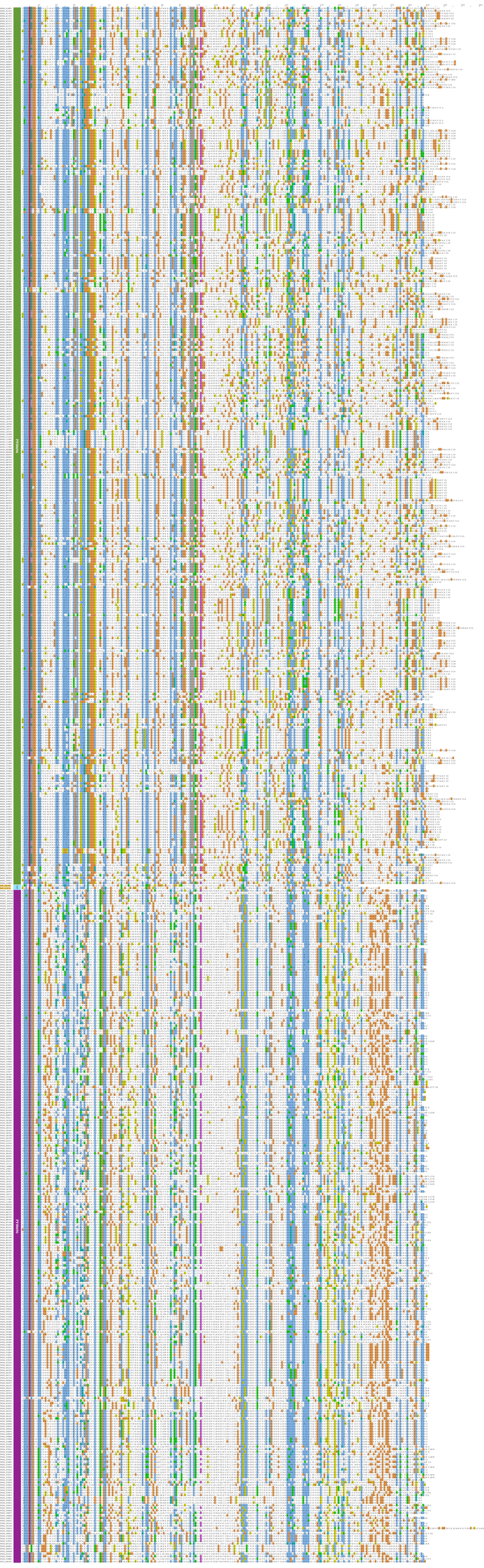
