## Supplementary Data 1 for "Deep embeddings to comprehend and visualize microbiome protein space"

**Supplementary Data 1.** Domain sequence alignment between two outlying proteins (BUK_OCEIH and BUK_DEIRA) and other butyrate kinase proteins (EC 2.7.2.7)

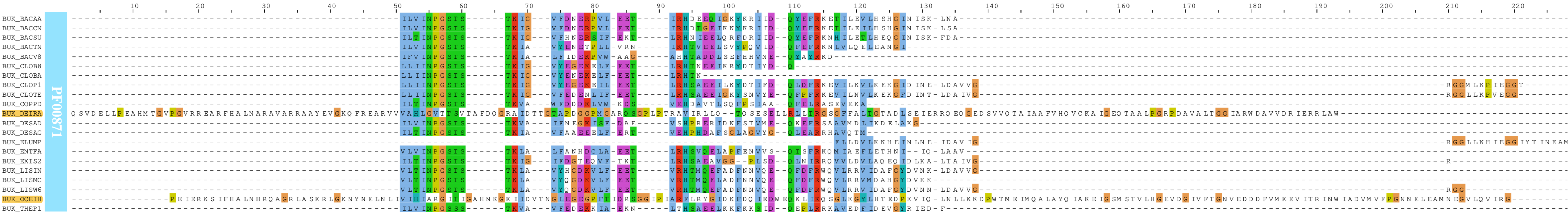
