## Supplementary Data 3 for "Deep embeddings to comprehend and visualize microbiome protein space"

**Supplementary data 3.** Domain sequence alignment between two outlying proteins (BUK_OCEIH and BUK_DEIRA) and PF00871 domain sequences from other phosphotransferases (EC 2.7.2.1, 2.7.2.15 and 2.7.2.7)

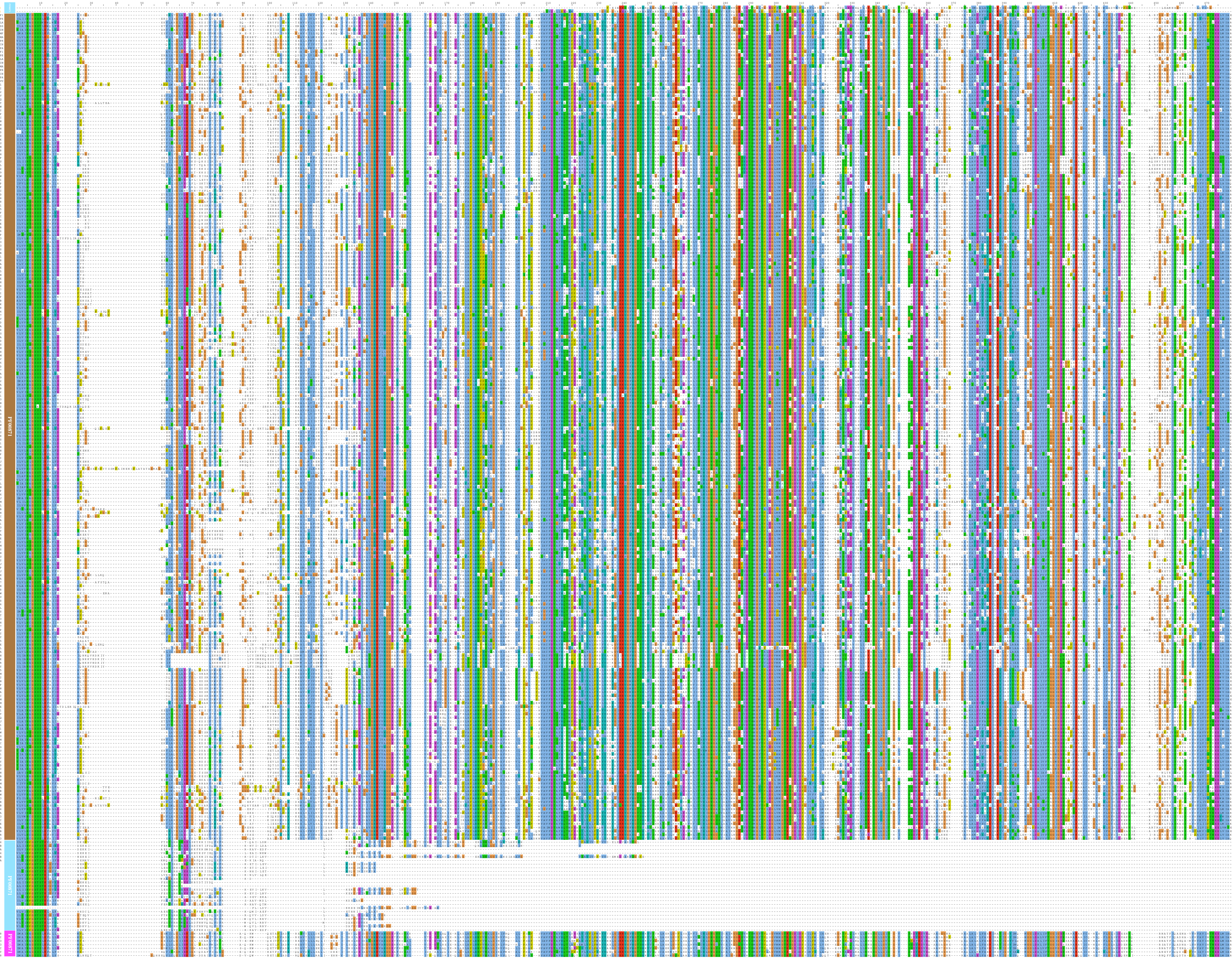
